# High-dimensional HIV-1 quasispecies modeling guides escape-proof antibody design

**DOI:** 10.64898/2026.09.02.748882

**Authors:** Griffin Kutler Dodd, Paulien Hogeweg, Rob J. de Boer

## Abstract

Rapidly evolving viruses form diverse quasispecies that enable escape from immune responses and treatments. For example, HIV-1 can rebound within weeks of broadly neutralizing antibody (bNAb) treatment through the outgrowth of high-fitness escape mutants in the quasispecies or the evolution of new escape variants. Most existing models of viral dynamics consider only a small number of viral variants and either assume arbitrary mutant fitness distributions or require extensive fitting to sparse clinical data. Here, we develop a high-dimensional HIV-1 quasispecies model that captures the dynamics of millions of viral strains and parameterize this using *in silico* binding affinity predictions. Without fitting to experimental data, the model qualitatively reproduces viral rebound following bNAb treatment. Lower-dimensional model projections recover these dynamics only when informed by features derived from the high-dimensional model. Finally, we use the model to develop a quasispecies-based framework for antibody optimization and identify antibodies predicted to effectively suppress viremia. Together, our results demonstrate that integrating mechanistic genotype-phenotype maps with high-dimensional quasispecies models provides unprecedented insights into viral evolution.

## Introduction

The high mutation rates of RNA viruses enable them to rapidly escape from host immune responses and from treatments. The diversity of rapidly evolving viruses within a host population means they are best thought of as broad clouds of related mutants, termed quasispecies (*1, 2*). For viruses such as influenza and SARS-CoV-2, escape occurs at the population level, with new vaccines needing to be developed to keep pace with rapid antigenic evolution in the global quasispecies (*3, 4*). Strikingly, the quasispecies of some viruses are so broad that escape can readily occur within an infected host. The combination of high mutation and viral load during HIV-1 infection means every possible single-base mutation is expected to occur each day (*5*), allowing HIV-1 to continually escape antibody and CD8^+^ T cell responses throughout infection (*6, 7*).

Crucially, variants present at very low frequencies in the HIV-1 quasispecies have been shown to be important drivers of immune escape (*8*) and treatment resistance (*9, 10*), often at frequencies that are inaccessible to most untargeted sequencing approaches (*11*). For example, variants resistant to the broadly neutralizing antibody (bNAb) 10-1074 were estimated to be present at 0.01%-0.1% frequency before administration and led to complete rebound of the viral load within weeks of treatment (*12, 13*). Understanding HIV-1 evolutionary dynamics in response to treatment therefore requires computational approaches capable of accounting for quasispecies diversity.

Most existing models of HIV-1 dynamics rely on clinical data to estimate the fitness and initial frequencies of commonly observed escape mutants (*13–16*). However, these models require extensive clinical data and provide limited mechanistic insight into the evolutionary dynamics driving viral adaptation. The attempts made at HIV-1 quasispecies models (*17–20*) assume a fixed distribution of mutant fitnesses, thereby not capturing the effect of a realistic genotype-phenotype mapping on quasispecies evolution. To overcome these limitations, here we introduce a novel simulation framework leveraging *in silico* binding affinity predictions to study viral quasispecies dynamics at an unprecedented scale. We apply this framework to study the evolution of millions of HIV-1 variants, both during untreated infection and after bNAb treatment.

## Results

### Quasispecies evolution on a receptor-binding landscape

Our model is based on a classical within-host ordinary differential equation model of HIV-1 dynamics, and is extended to accommodate stochastic mutations between strains. We first computed a realistic genotype-phenotype map by deriving a mechanistic relationship between receptor-binding affinity and viral infection rate (see Methods). We focus on the evolution of five non-glycosylated amino acids in the CCR5-binding site of Env that have previously been associated with escape from the broadly neutralizing antibody 10-1074 (*21*). Considering all amino acid combinations at these five residues yields a genotype space of 3.2 million virus variants. For each variant, we predicted the binding affinity to the CD4/CCR5 receptor complex using EvoEF, a physical energy function for estimating the effect of mutations on protein-protein binding affinity (*22*).

Before simulating quasispecies evolution, we first characterized the topography of the receptor-binding landscape. We used the reference strain 92BR020 as a wild-type sequence because a crystal structure is available for its Env protein in complex with CD4/CCR5. Among the 3.2 million viral variants, only 0.7% were predicted to bind the CD4/CCR5 complex more strongly than 92BR020 (Fig. 1A). 536 variants were local fitness peaks where no single mutation could further increase receptor-binding affinity. This low number of fitness peaks is strikingly different compared with the ruggedness observed in experimentally measured protein fitness landscapes (*23, 24*) and theoretical landscape distributions (Supplementary Text), revealing an exceptionally smooth receptor-binding landscape at these residues. Moreover, the fitness peaks are much closer together in the landscape than random sequences (Fig. 1B), indicating that the highest-affinity variants occupy a restricted region of sequence space. Network analysis showed that the 536 peak genotypes formed just four connected components (neutral ridges), each comprising variants with identical receptor-binding affinity, thereby forming neutral networks (Fig. 1C). Together, the smoothness and extensive neutrality of the landscape suggest that the quasispecies can readily evolve towards the fittest variants (*25, 26*).

**Figure 1:**
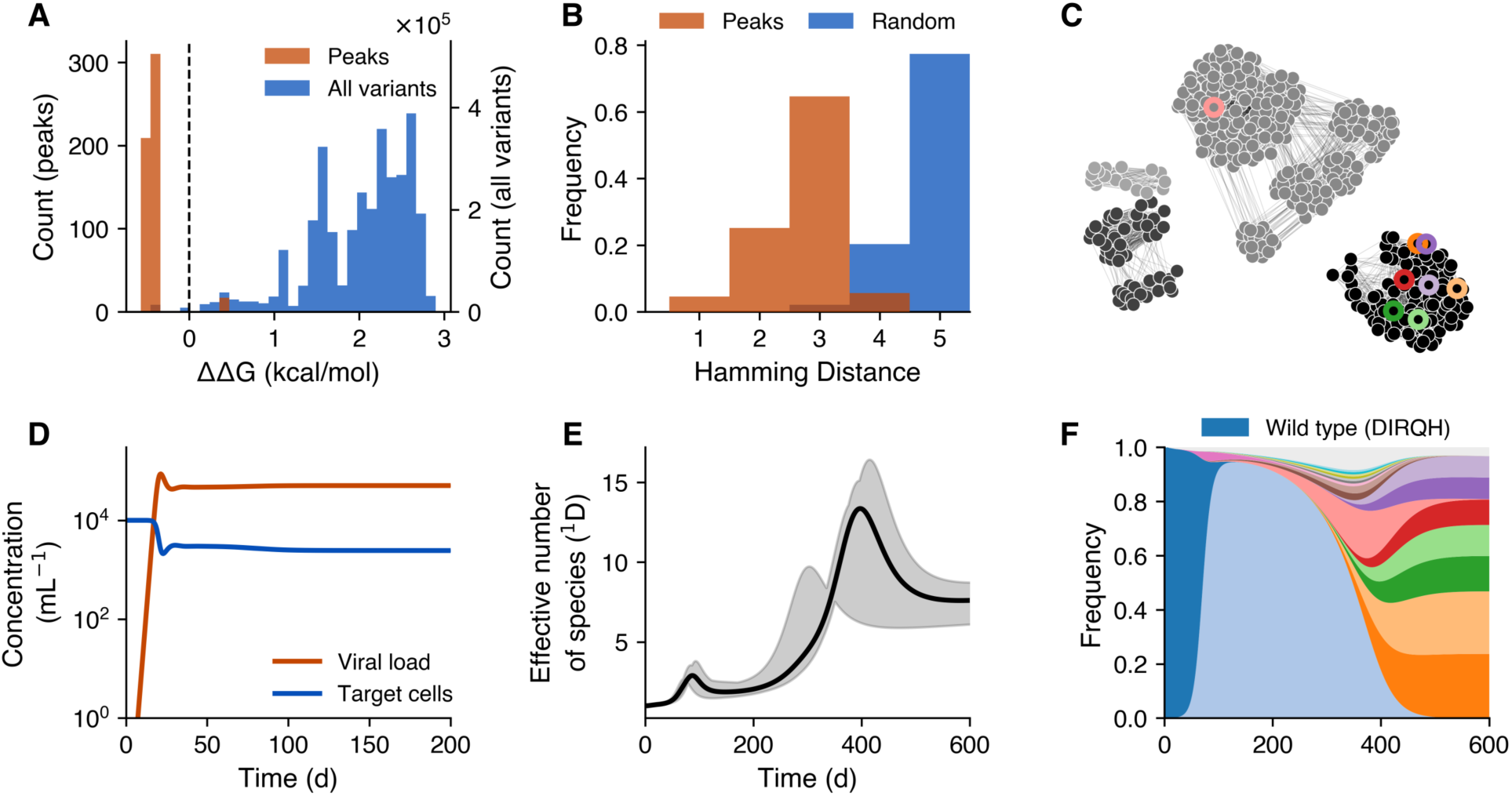
The HIV-1 quasispecies readily navigates a receptor-binding landscape. **(A)** Distribution of predicted binding affinities to the CD4/CCR5 complex for 3.2 million viral Env variants (blue) and for the 536 local fitness peaks (orange), shown relative to the clade B isolate 92BR020 Env (dashed black line). **(B)** Distribution of pairwise sequence distances between local fitness peaks (orange) compared with 536 random sequences (blue). **(C)** Network representation of fitness peaks, with edges connecting variants separated by a single mutation. Nodes are colored by receptor-binding affinity, with darker colors indicating higher affinity. **(D)** Viral load (orange) and target cell counts (blue) from ten simulated infections initiated with 92BR020. **(E)** Quasispecies diversity over time, measured as the effective number of species (the exponential of Shannon diversity, ^1^D). The solid line shows the mean across simulations, and the shaded region indicates the minimum and maximum values. **(F)** Frequencies of the 20 most abundant viral variants over time from a representative simulation, with each sequence shown in a different color. Nodes in (C) are outlined with their corresponding color from (F).

To test this prediction, we simulated quasispecies evolution, starting the infection with 92BR020 as a founder variant. As in classical target cell-limited models, viral load increased exponentially before contracting to a stable set point (Fig. 1D). Quasispecies diversity was highly dynamic, remaining low during infection, increasing rapidly after the set point was reached, and then declining as the quasispecies approached eco-evolutionary equilibrium (Fig. 1E). Notably, the expansion in diversity followed by contraction resembles previously reported patterns of within-host HIV-1 diversity (*27–29*), although the timescale in our model is much faster since we consider evolution at only five residues. Tracking the most abundant variants revealed that diversity peaked as the quasispecies approached the highest binding affinity neutral network and declined as the population became localized on that network (Fig. 1F). Evolutionary trajectories were reproducible across simulations, differing primarily in the final proportions of variants on the neutral network (Fig. S1). There was neutral drift after the quasispecies localized on this ridge, but this occurred on a long timescale. Together, these results establish a framework for tracking quasispecies evolution across millions of viral genotypes.

### High-dimensional modeling reproduces antibody responses

We next used our model to study how an evolving quasispecies responds to broadly neutralizing antibody treatment. We predicted the binding affinity of the same 3.2 million virus variants to the bNAb 10-1074 using EvoEF, and fit a biphasic pharmacokinetic model to serum antibody concentrations measured in a clinical trial to capture antibody decay (Fig. S2) (*12*). Simulated treatment with 30 mg/kg of 10-1074, the highest dose used in clinical trial, led to viral rebound regardless of antibody administration time (Fig. 2A and 2B). Consistent with clinical observations, rebound occurred on the timescale of weeks across all simulations. Rebound was fastest when treatment was administered near the peak in quasispecies diversity, approximately one year after infection, and treatment at one and two years selected for genetically different escape mutants (compare Fig. 2C and 2D). We also observed a transient overshoot in viral load before contraction to the pre-treatment set point. This overshoot was observed in some patients after 10-1074 treatment (*12*), although sampling near this peak was often sparse. Notably, a previous low-dimensional model fit to the same clinical data did not produce this overshoot (*13*), whereas our model reproduces this feature without being fit to the data.

**Figure 2:**
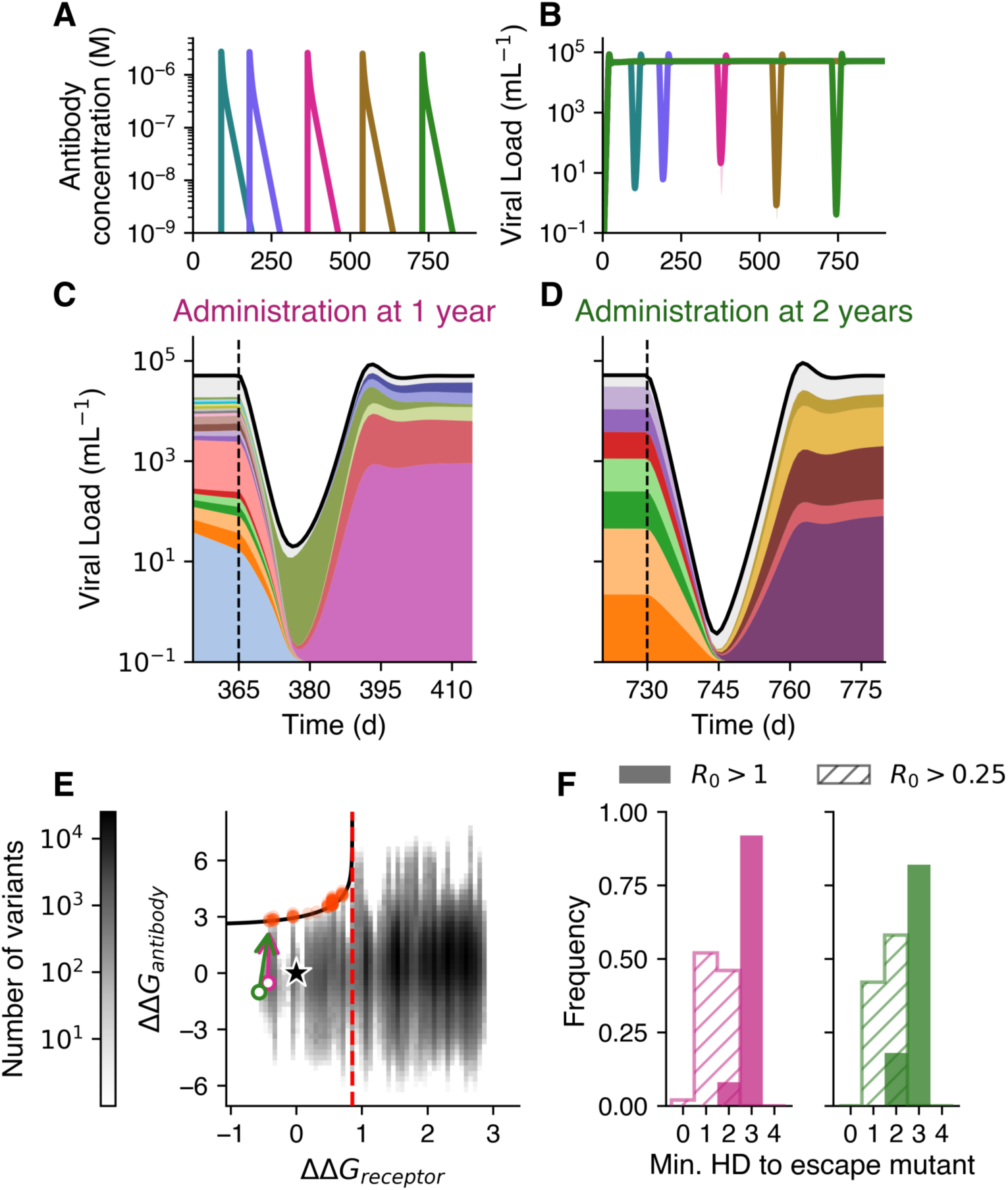
Quasispecies modeling reproduces viral rebound after administration of the broadly neutralizing antibody 10-1074. **(A and B)** Simulated treatment with 30 mg/kg of 10-1074 administered at five different times post-infection. Antibody decay kinetics (A) were fit to patient serum antibody concentrations, and viral loads (B) are shown as the mean (thick line) and minimum/maximum (lightly shaded region) across ten simulations for each treatment time. **(C and D)** Frequencies of the most abundant variants following administration (C) one year or (D) two years post-infection. Colors of the 20 most abundant variants before treatment match Fig. 1F, and the ten most abundant variants after treatment are assigned new colors. **(E)** Distribution of receptor- and antibody-binding affinities for all 3.2 million viral sequences, shown relative to 92BR020 Env binding to CD4-CCR5 (ΔΔ*G_receptor_*) and 10-1074 (ΔΔ*G_antibody_*); 92BR020 is marked with a star. The dashed red line and black contour indicate *R*_0_ = 1 in the absence of antibody and at an antibody concentration of 30 mg/kg, respectively. Colored arrows trace the mean receptor- and antibody-binding affinities of the quasispecies from the time of treatment to 50 days after treatment in the simulations shown in (C) and (D) (pink and green, respectively). **(F)** Minimum Hamming distance between the 50 most frequent variants at the time of treatment administration and variants with *R*_0_ *>* 1 (solid) or *R*_0_ *>* 0.25 (hatched). Pink and green histograms indicate treatment administered one and two years post-infection, respectively.

To identify which variants drive viral rebound after antibody administration, we derived an expression for the basic reproduction number *R*_0_ of each variant as a function of receptor-binding affinity, antibody-binding affinity, and antibody concentration (Supplementary Text). Only a small subset of variants had *R*_0_ *>* 1 immediately following treatment with 30 mg/kg of 10-1074 (red points in Fig. 2E). Because *R*_0_ depends on antibody concentration, decaying antibody concentration progressively expands the set of variants capable of sustaining infection. We therefore asked whether rebound results from rapid localization on initially viable escape mutants. Although there was selection for lower antibody-binding affinity in the quasispecies after treatment, the mean was still *R*_0_ *<* 1 at the maximal dose of antibody after treatment (arrows in Fig. 2E). Consistent with this, maintaining antibody concentration at 30 mg/kg always resulted in viral extinction, whereas reducing the concentration tenfold enabled viral rebound (Fig. S3). Moreover, the 50 most abundant variants present at one year were at least two mutations away from any variant with *R*_0_ *>* 1, whereas variants with *R*_0_ *>* 0.25 at maximal antibody dose (which will later become *R*_0_ *>* 1 when the antibody concentration declines) were accessible by a single mutation (Fig. 2F). Together, these results show that viral rebound depends on antibody decay that progressively expands the set of viable escape variants.

Having shown that antibody escape emerges naturally from a high-dimensional quasispecies model, we asked whether these dynamics could be captured in a lower-dimensional projection. We considered only features measurable before antibody treatment, enabling dimensionality reduction without fitting to bNAb clinical trial data. We first selected variants using experimentally accessible low-dimensional features: viral fitness and frequency. Ranking variants by their fitness in the absence of antibody required nearly 10^4^ strains to consistently produce rebound across simulations with different treatment administration times (Fig. 3A). Adding all variants within one mutation of the founder improved performance, likely by restoring mutational paths between the founder and the highest fitness variants, but these projections still required at least 96 variants (5×19+WT) and reproduced rebound in fewer than half of simulations. Frequency-based selection performed substantially better: including all variants with a maximum frequency above 1% at any point during the simulation yielded a model with 28 variants that consistently reproduced rebound. However, frequency selection requires longitudinal measurements of quasispecies composition, and a 1% frequency is approaching the detection limit of next-generation sequencing (*11*). Moreover, in the full HIV-1 sequence space, this threshold would likely include many more variants, resulting in a substantially higher-dimensional model.

**Figure 3:**
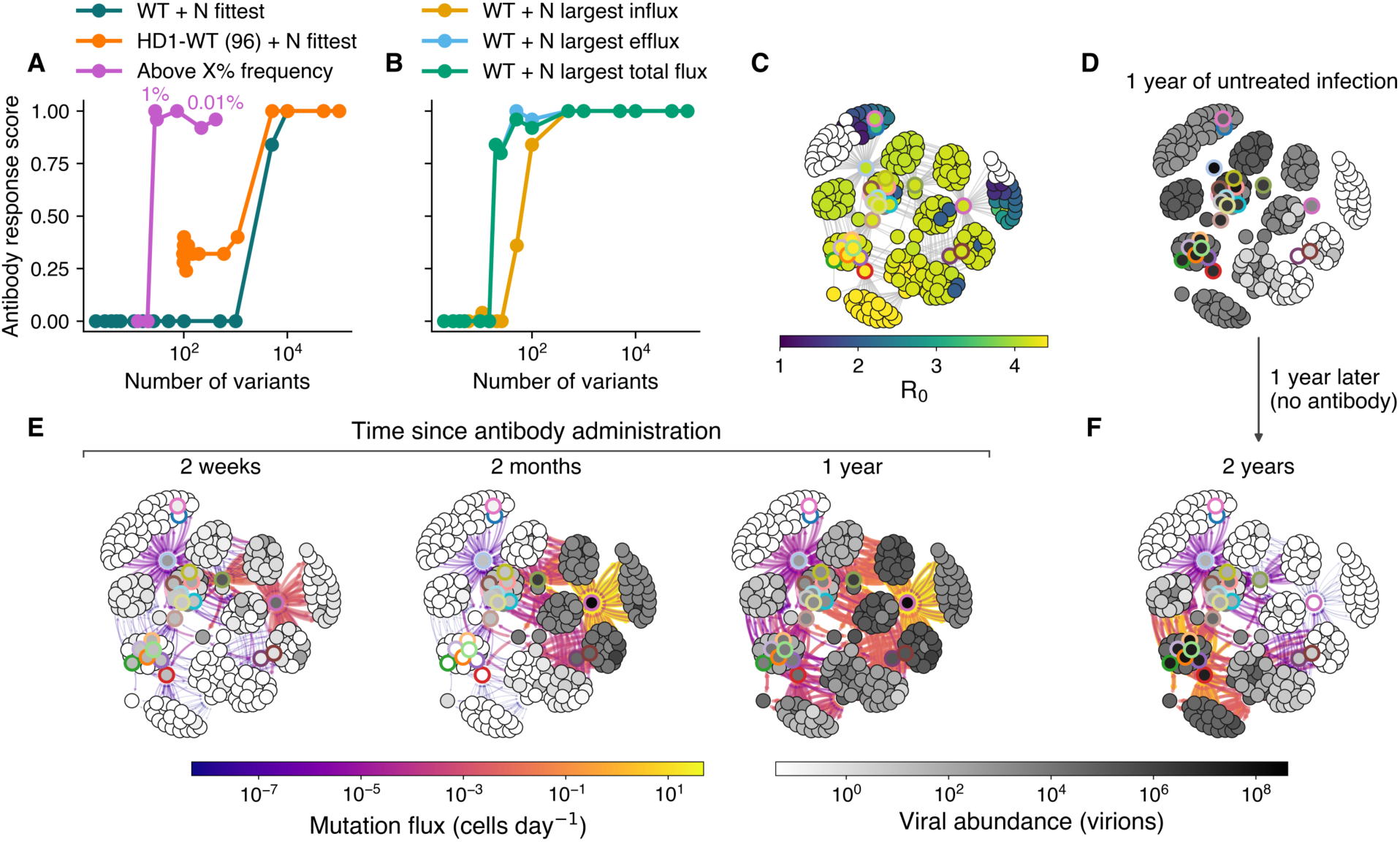
High-dimensional quasispecies features identify low-dimensional projections that capture antibody rebound. **(A and B)** Performance of low-dimensional model projections in reproducing antibody rebound. The antibody response score is the proportion of 25 simulations (five at each of five antibody administration times) where rebound occurs. Variants were selected using (A) experimentally accessible features measured in the absence of antibody: viral fitness, Hamming distance from the founder variant, and maximum frequency at any timepoint in simulated untreated infections; or (B) features derived from the high-dimensional quasispecies: total mutational influx (cumulative inbound mutations), total mutational efflux (cumulative outbound mutations), or total mutational flux (the sum of influx and efflux). **(C-F)** Graph visualization of the quasispecies from a simulated infection treated with 10-1074 after one year. Nodes represent viral variants, edges connect variants separated by a single mutation, and only variants with the highest cumulative mutational fluxes are shown. Nodes are colored by (C) fitness (*R*_0_ in the absence of antibody, with *R*_0_ *<* 1 variants shown in white), (D) abundance immediately before antibody administration, (E) abundance at successive times following treatment, and (F) abundance one year later in a matched untreated simulation (two years post-infection). In (E) and (F), edge color indicates the mutational flux between connected variants. Nodes are outlined with their corresponding color in Fig. 2B and C, and panels (D-F) are stills from movie S1.

We next tested whether the high-dimensional model itself could identify informative low-dimensional projections. To this end, we quantified the mutational flux between neighboring variants as the time-integrated abundance of cells infected by a source variant multiplied by the mutation probability to the neighboring genotype. Selecting variants with the largest total mutational reproduced rebound in more than 80% of simulations using only 20 variants (Fig. 3B). Rankings based on the total mutational influx or total flux (summed influx and efflux) were slightly less successful compared with efflux, but all three approaches were capable of consistently reproducing rebound using on the order of 100 variants. Together, these results show that low-dimensional models can capture complex dynamics like antibody rebound, but only when informed by deep sequencing data or features extracted from high-dimensional models.

To understand why many low-dimensional projections failed to capture antibody rebound, we explored how antibody treatment reorganizes quasispecies composition. We constructed a graph in which nodes represent virus variants, edges connect variants separated by a single mutation, and we depict only the variants with the largest mutational fluxes in the full-dimensional simulation. Although most included variants had high fitness, the graph also contained several clusters composed predominantly of variants with *R*_0_ *<* 1 (Fig. 3C). Before antibody administration, the quasispecies was concentrated on a subset of high-fitness variants (Fig. 3D) corresponding to part of the highest-fitness neutral network in Fig. 1C. Following treatment, the quasispecies rapidly redistributed into previously sparsely occupied clusters of lower-fitness variants (Fig. 3E). Remarkably, this treatment-induced reorganization persisted for more than a year after antibody administration, remaining markedly different from that of an untreated infection of the same duration despite the antibody long having decayed (Fig. 3F). As a consequence, repeated antibody administration was less effective, with second doses exhibiting reduced efficacy even when administered months after the initial treatment (Fig. S4). Thus, antibody treatment can reshape the quasispecies in ways that alter responses to subsequent treatment long after the initial dose.

### Antibody design accounting for viral diversity

bNAbs are typically evaluated by their neutralization potency against reference virus panels. Although these panels are designed to capture viral diversity, they typically contain only tens to hundreds of variants (*30*), representing a substantial reduction in dimensionality compared to the natural diversity during infection. We used our high-dimensional model to ask how different approaches to this dimensionality reduction affect the performance of designed bNAbs. Starting from wild type 10-1074, we generated all 95 single amino acid substitutions at five residues near the virus-binding interface and used EvoEF to predict the binding affinity of each candidate antibody to every virus in a given panel. We then selected the antibody that minimized the maximum *R*_0_ among viruses in the panel in the presence of the antibody. Antibody selection used an *ɛ*-greedy scheme, where the best antibody was selected with probability 1 − *ɛ* and a random candidate with probability *ɛ*, with *ɛ* decreasing exponentially each iteration. From the selected antibody, we again made the all 95 single substitutions at the same five residues and repeated the process until convergence (Fig. 4A).

**Figure 4:**
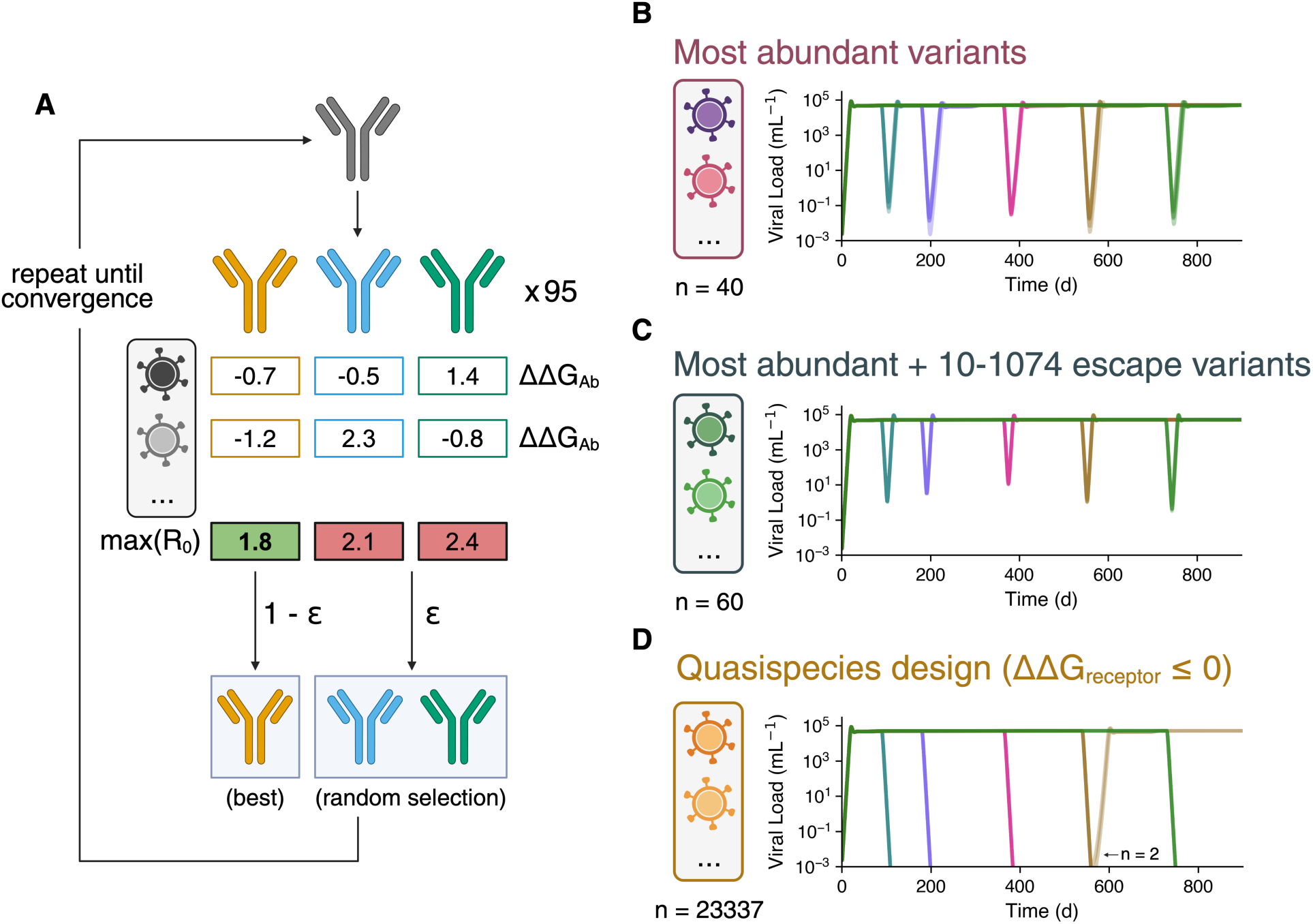
Quasispecies design finds effective broadly neutralizing HIV-1 antibodies. **(A)** Antibody optimization workflow. Starting from a given antibody, all single amino acid substitutions at five selected residues are generated. For each candidate antibody, EvoEF is used to predict binding affinities to a panel of target viruses which, together with their receptor-binding affinities, are used to calculate the *R*_0_ of each virus in the presence of the antibody. At each iteration, the candidate that minimizes the maximum *R*_0_ across the virus panel is selected with probability 1 − *ɛ*, while a randomly selected candidate is chosen with probability *ɛ*, with iterations proceeding until convergence. **(B-D)** Viral loads following simulated treatment with 30 mg/kg of the best antibody (lowest maximal *R*_0_) identified across 25 optimization runs against virus panels comprising (B) the 40 most abundant variants during untreated infection (Fig. S1); (C) these 40 variants plus the 20 most abundant variants following antibody administration; or (D) all variants with receptor-binding affinity at least as strong as the wild type (*quasispecies design*). Created in https://BioRender.com.

We first tested a panel comprising the 40 most abundant variants during untreated infection, mimicking panel designs intended to capture natural diversity. Optimizing 10-1074 binding against these variants improved viral suppression following treatment, but the resulting antibody was still unable to prevent rebound (Fig. 4B). We next expanded the panel with the 20 most abundant variants following 10-1074 treatment (Fig. 2B and 2C). Optimization against this 60-variant panel identified an antibody only one mutation away from wild type 10-1074 that completely suppressed viremia in our simulations (Fig. S5); notably, none of the 3.2 million variants in our model had *R*_0_ *>* 1 in the presence of 30 mg/kg of this antibody (Fig. S6). However, the global optimized antibody identified using this panel was even less effective than wild type 10-1074 (Fig. 4C), suggesting that optimization had overfit to the variants in the panel. Indeed, although this optimized antibody strongly bound the high-frequency variants in the quasispecies, there were now many new escape variants with *R*_0_ *>* 1 (Fig. S6). This decrease in performance highlights that effective neutralization of a low-dimensional reference virus panel does not necessarily predict antibody performance.

We next relaxed the dimensionality reduction by constructing a panel comprising all variants with receptor-binding affinity at least as strong as the reference wild type (*n* = 23, 337), which we term *quasispecies design* because the panel begins to approach the diversity of the full quasispecies. Optimization against this panel identified the same effective antibody one substitution from wild-type 10-1074 as the 60-variant panel (Fig. S5), while the global optimum antibody suppressed rebound in 23/25 simulations (Fig. 4D). Thus, quasispecies design identified multiple effective bNAbs without prior knowledge of escape variants from the wild-type antibody. We then asked whether mutational efflux, which was the most effective high-dimensional feature for constructing model projection, could also be used to design an effective low-dimensional panel. However, a panel comprising the 60 variants with the highest efflux during untreated infection identified the same antibodies as the panel with the 40 most abundant variants (Fig. S5), suggesting features that capture quasispecies dynamics in reduced models are not necessarily good features for designing reference panels for antibody optimization. Together, our results demonstrate that model dimensionality strongly affects treatment design and predicted outcomes, highlighting the value of our high-dimensional approach for capturing dynamics that may be missed by lower-dimensional models.

## Discussion

Here, we developed a high-dimensional within-host model that couples a mechanistic genotype-phenotype map to viral evolution across millions of variants. While previous models have incorporated high viral diversity using arbitrarily specified fitness landscapes (*19, 31*), we show that the structure of a natural genotype-phenotype mapping can be exploited for treatment design, and that effective treatments are only found when the high-dimensionality of the landscape is retained. A similar approach was recently developed by Mohanty & Shakhnovich, who used EvoEF to compute high-dimensional receptor- and antibody-binding landscapes for SARS-CoV-2 and showed how these landscapes could guide vaccine design (*32*). However, their evolutionary simulations have several limitations: they considered viral diversity only at the between-host level, limited the number of variants present at any given time, and assumed constant antibody concentrations. We extended this framework to within-host HIV-1 evolution without restricting the number of possible variants and incorporated antibody pharmacokinetics, making viral fitness time-dependent. Our model revealed that highly diverse quasispecies can explore evolutionary paths to variants that are unfit at peak antibody concentrations but become viable as antibody levels decline, enabling viral rebound from initially effective antibody treatment.

Our model makes several simplifying assumptions that are relevant to address in future work. We assume that all virally infected cells are short-lived and therefore do not model the latent reservoir. Reactivation of long-lived latently infected cells could make single-dose antibody treatment more difficult, as their lifespan during antiretroviral therapy is far longer than the duration of therapeutic antibody concentrations (*33*). Incorporating latency would also introduce evolutionary memory, as reactivation can reintroduce previously abundant variants into the quasispecies. Consistent with this, Doekes et al. have previously shown that reactivation of less infectious viral strains from the latent reservoir can slow the evolution of virulence (*20*). We also do not model immune responses against infected cells. A subset of patients durably control viral rebound following bNAb administration, and this post-treatment control is associated with faster CD8^+^ T cell proliferation (*34, 35*). Previous models have shown that post-treatment control can be explained by bistability, in which CD8^+^ T cells suppress reactivation of latently infected cells under some parameter regimes (*36–38*). Extending our model to include latency and immune responses could reveal how the interplay of viral diversity and host immunity determines the success of antibody treatment.

Beyond these biological extensions, the rapidly expanding repertoire of accurate and fast machine learning methods for predicting viral phenotypes provides new opportunities to expand the genotype-phenotype maps underlying mechanistic models. Tools are now available for predicting protein binding affinities and stability (*39–42*), antigenic distances and cross-reactivity (*43–45*), host tropism and receptor usage across diverse viruses (*46–48*), and a growing range of other viral phenotypes. We chose EvoEF for its computational efficiency, but our framework could readily incorporate new methods as they become available. Integrating predictive tools with mechanistic models will enable viral evolution to be studied at scales far beyond those accessible to experiments. Our work represents a step towards this integration, showing how computational prediction of genotype-phenotype maps can transform mechanistic models from tracking a small number of variants into models of entire viral quasispecies.

## Methods

### Residue selection and binding aAnity predictions

Viral escape from the CCR5-binding broadly neutralizing antibody 10-1074 has been attributed to a small number of mutations in the HIV-1 Env protein (*12, 21, 49*). These mutations either disrupt glycosylation or alter amino acids within the CCR5-binding site. Because current structure-based energy functions cannot reliably predict the effect of changes in glycosylation on binding affinity, we restricted our analysis to five non-glycosylated residues in the V3 loop of gp120 (D325, I326, R327, Q328, H330).

Virus-receptor and virus-antibody binding affinities were predicted using EvoEF, a physics-based energy function for estimating the effect of mutations on protein-protein binding affinity (*22, 32*). Receptor-binding affinities were predicted using the crystal structure of gp120 from the reference strain 92BR020 in complex with CD4 and CCR5 (PDB 6MEO). Antibody-binding affinities were predicted using the structure of the BG505 SOSIP Env trimer in complex with the broadly neutralizing antibodies 10-1074 and BG24 (PDB 7UCF), retaining only the gp120-10-1074 complex (chains G, H, and L). We predicted changes in receptor- and antibody-binding affinities ΔΔ*G* for the combinatorially complete set of viral mutants at these residues (3.2 x 10^6^ *–* 1 mutants).

### Quasispecies model

We began with the classical two-stage within-host HIV-1 model, which describes the dynamics of susceptible target cells, infected cells, and free virus (Equations 1-4). Target cells *T* are lost through death or deactivation at rate *d* and become infected by a single viral strain *V* at rate *β*, producing eclipse-phase *I*_1_ infected cells. *I*_1_ cells either die at rate *δ*_1_ or mature into virus-producing *I*_2_ cells at rate γ (meaning we neglect superinfection). Virus-producing cells die at rate *δ*_2_ and produce *n* virions during their lifetime, which are cleared at rate *c* (Equation 4).

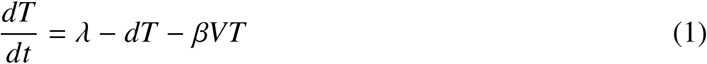

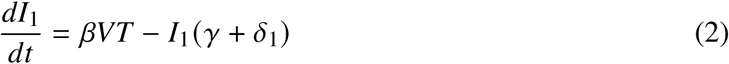

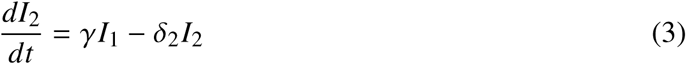

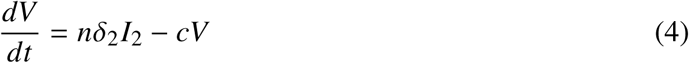

We extended this model in three ways to allow us to study quasispecies dynamics with our computed genotype-phenotype map. First, we derived a mechanistic relationship between receptor-binding affinity and the infection rate *β* (Supplementary Text), such that the infection rate of a mutant *i*, *β_i_*, depends on the wild-type infection rate *β_wt_* and the change in receptor-binding affinity ΔΔ*G_i_*, according to

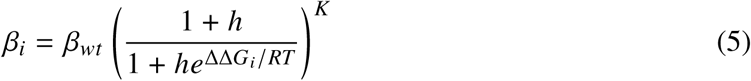

where *RT* is product of the ideal gas constant and temperature, and *h* and *K* are saturation and shape parameters that determine the maximum fitness gain relative to the wild type.

Second, we coupled viral strains through mutation by defining a mutation matrix *μ*, where element *i, j* denotes the probability that strain *i* mutates to strain *j*, and *μ_ii_* is the probability that no mutation occurs. Because HIV-1 mutations occur primarily during genome integration (*50*), we modeled mutation during the transition from *I*_1_ to *I*_2_, with each *I*_2_ cell subsequently producing the same virus variant.

Finally, we incorporated antibody dynamics by deriving the equilibrium fraction of free (unbound) virions of strain *i*, *θ_i_*, for each strain:

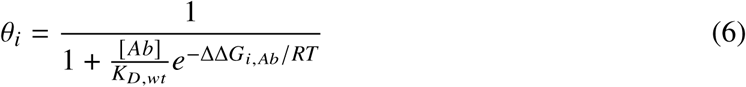

which depends on the antibody concentration [ *Ab*], the dissociation constant of the wild type virus from the antibody *K_D,wt_*, and the change in antibody-binding affinity ΔΔ*G_i,Ab_* (Supplementary Text). Because only free virions can infect target cells, the effective infection rate of strain *i* is given by *θ_i_ β_i_*. The complete model is therefore

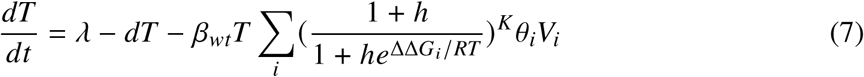

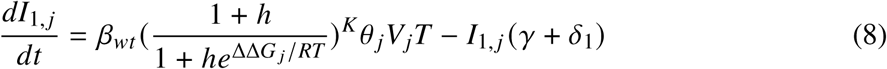

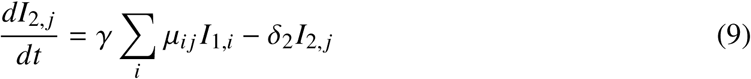

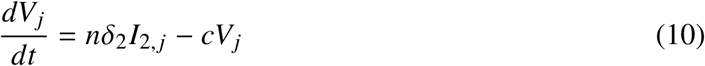

The model was simulated using Euler integration with a step size of Δ*t* = 0.01 d. Because the full mutation matrix *μ* contains 3.2 x 10^6^ x 3.2 x 10^6^ entries, we represented it implicitly as the Kronecker product of five 20×20 per-site mutation matrices (*51, 52*). To prevent instant mutational spreading of the quasispecies, mutations with an expected occurrence of fewer than one per time step were treated probabilistically, with the probability of occurrence equal to the expected mutational flux. All model parameters are given in Table S1.

### Pharmacokinetic modeling and antibody dosing

Passive infusion of broadly neutralizing antibodies produces biphasic decay kinetics (*53*). We therefore modeled antibody concentration [ *Ab*](*t*) following completion of infusion as:

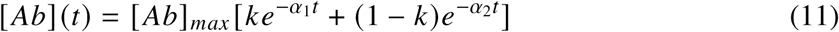

where [ *Ab*]*_max_* is the initial antibody concentration, *k* determines the relative contributions of the two decay phases, and *α*_1_ and *α*_2_ are the corresponding decay rates.

This model was fit to serum antibody concentrations from all patients given 30 mg/kg of 10-1074 in the Caskey et al. clinical trial (*12*). For each patient, the day 0 measurement was excluded and the parameters [ *Ab*]*_max_*, *k*, *α*_1_, and *α*_2_ were fit in log-space using nonlinear least squares (Fig. S2). Two patients with extreme fitted decay rates (1HD6K and 1HC2) were excluded, and the mean values of *k*, *α*_1_, and *α*_2_ across the remaining patients were used for the simulations.

To simulate treatment, we set [ *Ab*]*_max_* to the expected peak concentration following a 30 mg/kg infusion. Estimating a blood volume of 5 L, a patient weight of 70 kg, and the molecular weight of 10-1074 to be 150 kDa yields [ *Ab*]*_mac_* = 2.8 x 10^−6^ M. Using a wild-type dissociation constant of *K_D_*= 10^−8^ M (*54*) yields a free-virus fraction for the wild-type strain of *θ* = 0.004 at the peak antibody concentration (i.e., 99.6% of wild type virus is bound by antibody immediately after infusion).

### Antibody optimization

For antibody optimization, we selected five residues at the 10-1074 virus-binding interface where single mutations significantly altered the predicted receptor-binding affinity: G106, F110, and E112 on the heavy chain and S96 and R97 on the light chain (PDB 7UCF). Starting from the wild type 10-1074 sequence, we generated all 95 possible single amino acid substitutions at these positions. For each mutant antibody, EvoEF was used to compute binding affinities against four panels of viral variants: 1) the 40 most abundant variants before administration (Fig. 1); 2) these 40 variants together with the 20 most abundant escape variants observed across all administration times and replicates (50 total simulations); 3) the 60 variants with the largest total mutational efflux in simulations without antibody; or 4) all variants with receptor-binding affinity at least as strong as the wild type (i.e., ΔΔ*G* ≤ 0, *n* = 23337).

Each candidate antibody was scored by the maximum *R*_0_ across the selected virus panel in the presence of 30 mg/kg of antibody (i.e., by the best escape variant). Antibody optimization was performed using an *ɛ*-greedy search (*55*). At each iteration, the candidate antibody with the lowest maximum *R*_0_ (i.e., the one with the lowest fitness escape variant) was selected with probability *ɛ*; otherwise, one of the 95 mutated antibodies was chosen at random. The exploration probability decayed exponentially according to *ɛ*_0_×exp[− *δ*×iteration], with*ɛ*_0_ = 0.3 and *δ* = 0.3. Optimization was terminated when the same antibody was selected in two consecutive iterations. We performed 25 independent optimization runs for each virus panel and evaluated both the best antibody identified after a single mutation and the overall optimized antibody with the lowest maximum *R*_0_ from each panel.

## Acknowledgments

We thank Bram van Dijk for useful advice on data visualization and Jan Kees van Amerongen for maintaining our computer cluster, which made these simulations possible.

## Author contributions

Conceptualization: all authors; Methodology: all authors; Formal analysis: G.K.D.; Software: G.K.D.; Visualization: G.K.D.; Writing - original draft: G.K.D.; Writing - review & editing: all authors.

## Data, code and materials availability

All code used to simulate the model and reproduce the figures is hosted on GitHub (https://github.com/gkutlerdodd/HIV-quasispecies-modeling). The simulation data and Movie S1 are available in a Zenodo repository (https://doi.org/10.5281/zenodo.22256270).

## Supplementary materials

Supplementary Text

Figs. S1 to S9

Table S1

References (*49–66*)

Movie S1

