## Supplementary Text, Figures S1-S9, Table S1 for "High-dimensional HIV-1 quasispecies modeling guides escape-proof antibody design"

### Supplementary Materials for
### High-dimensional HIV-1 quasispecies modeling guides escape-proof antibody design

Griffin Kutler Dodd*, Paulien Hogeweg, Rob J. de Boer

**This PDF file includes:**

Supplementary Text

Figures S1 to S9

Table S1

Captions for Movie S1

**Other Supplementary Materials for this manuscript:**

Movie S1

#### Supplementary Text

##### Relationship between receptor-binding affinity and infection rate

We derived a mechanistic relationship between receptor-binding affinity and the viral infection rate $\beta$ using the kinetics of virus-receptor binding. Let free virus $V$ bind receptors on target cells $T$ to form virus-receptor complexes $\hat{R}$ with association and dissociation rate constants $k_{on}$ and $k_{off}$, respectively. Viral entry into target cells occurs at rate $k_{entry}$. Considering that receptor occupancy is low (*56*) such that the number of free receptors is approximately the number of target cells multiplied by the number of receptors per cell $R$, the dynamics of virus-receptor complexes are given by:

$$\frac{d\hat{R}}{dt} = k_{on}RVT - k_{off}\hat{R} - k_{entry}\hat{R} \tag{S1}$$

Making a quasi-steady state assumption for virus-receptor complexes yields:

$$\hat{R} = k_{on}RVT/(k_{off} + k_{entry}) \tag{S2}$$

Because new infected cells ($I_1$) are generated by viral entry at rate $k_{entry}\hat{R}$, the influx of $I_1$ cells is $k_{on}k_{entry}RVT/(k_{off} + k_{entry})$. The short lifetime of gp120-CD4 complexes (*57*) suggests that dissociation is faster than viral entry cell entry ($k_{off}/k_{entry} >> 1$). Thus, using $K_D = k_{off}/k_{on}$, the influx of $I_1$ cells simplifies to $(k_{entry}R/K_D)VT$. Assuming $k_{entry}$ is constant between variants, the infection rate $\beta$ is inversely proportional to the receptor-binding dissociation constant $K_D$.

The infection rate of a mutant $\beta_i$ can therefore be expressed in terms of its change in receptor-binding free energy relative to the wild type $\Delta\Delta G_i$, as

$$\beta_i = \beta_{wt}e^{-\Delta\Delta G_i/RT} \tag{S3}$$

Because this exponential relationship predicts unrealistically large increases in $R_0$ for modest improvements in receptor-binding affinity[1], we introduced a saturation of the fitness benefit from increased receptor-binding affinity. We chose the Hill-like relationship:

$$\beta_i = \beta_{wt}(1+h)^K \left(\frac{e^{-\Delta\Delta G_i/RT}}{e^{-\Delta\Delta G_i/RT} + h}\right)^K = \beta_{wt}\left(\frac{1+h}{1+he^{\Delta\Delta G_i/RT}}\right)^K \tag{S4}$$

[1]For example, if the wild-type virus has $R_0 = 3.3$, a mutant with $\Delta\Delta G = -1$ kcal/mol would have $R_0 = 16.9$.

where $\beta_i = \beta_{wt}$ when $\Delta\Delta G_i = 0$. Model results were relatively robust to the exact form of the infection rate: simulations using the exponential relationship reproduced viral rebound following 10-1074 treatment and the performance of designed antibodies (Fig. S7).

To parameterize $\beta_{wt}$, we used the fact that the acute-phase growth rate of HIV-1 is between $\rho = 1$ d$^{-1}$ and 1.5 d$^{-1}$ (*58, 59*). $\beta$ in such a two-stage infection model has previously been solved as a function of $\rho$ (*60*):

$$\beta = \frac{cd(\delta_1 + \gamma + \rho)(\delta_2 + \rho)}{\lambda n \delta_2 \gamma} \tag{S5}$$

With our parameters (Table S1), and assuming $\rho = 1$ d$^{-1}$, we obtained $\beta_{wt} = 2\times10^{-9}$ virions$^{-1}$ d$^{-1}$.

**Antibody binding kinetics**

To model antibody neutralization, we considered the reversible binding of free virions ($V_i$) of strain $i$ to antibody (Ab), forming virus-antibody complexes $C_i$:

$$V_i + Ab \underset{k_{off,i}}{\overset{k_{on,i}}{\rightleftharpoons}} C_i \xrightarrow{\eta} \phi \tag{S6}$$

Assuming that virus-antibody complexes rapidly approach quasi-steady state,

$$C_i = \frac{k_{on,i}[Ab]}{k_{off,i} + \eta} V_i \tag{S7}$$

and the equilibrium fraction of free virions of strain $i$, $\theta_i$, is therefore

$$\theta_i = \frac{V_i}{V_i + C_i} = \frac{1}{1 + \frac{k_{on,i}}{k_{off,i}+\eta}[Ab]} \tag{S8}$$

When dissociation is much faster than complex clearance (i.e., $k_{off}/\eta >> 1$), this simplifies to:

$$\theta_i \approx \frac{K_{D,i}}{K_{D,i} + [Ab]} \tag{S9}$$

where $K_{D,i} = k_{off,i}/k_{on,i}$ is the equilibrium dissociation constant of strain $i$. Expressing $K_{D,i}$ in terms of the change in antibody-binding free energy $\Delta\Delta G_{i,Ab}$ relative to the wild-type virus yields:

$$\theta_i = \frac{1}{1 + \frac{[Ab]}{K_{D,WT}} e^{-\Delta\Delta G_{i,Ab}/RT}} \tag{S10}$$

#### $R_0$ calculation

We defined the fitness of a variant as the $R_0$ of that variant in the absence of mutations, which we computed using the next-generation matrix method (*61*). Let $\beta = \beta_{wt}(\frac{1+h}{1+he^{\Delta\Delta G_i/RT}})^K$. Considering the three infected compartments $I_1$, $I_2$, and $V$, and defining $F_i$ as the rate at which new infections appear in compartment $i$, the infection matrix $F$ is the Jacobian matrix of $F_i$ evaluated at the disease-free equilibrium $\bar{T} = \lambda/d$:

$$F = \begin{bmatrix} 0 & 0 & \frac{\beta\lambda\theta}{d} \\ 0 & 0 & 0 \\ 0 & 0 & 0 \end{bmatrix} \tag{S11}$$

Similarly, if $V_i$ is the outflow from compartment $i$ minus the inflow to compartment $i$ by other means, the transition matrix $V$ is the Jacobian of $V_i$ at the disease-free equilibrium:

$$V = \begin{bmatrix} \gamma + \delta_1 & 0 & 0 \\ -\gamma & \delta_2 & 0 \\ 0 & -n\delta_2 & c \end{bmatrix} \tag{S12}$$

The next-generation matrix $FV^{-1}$ is therefore:

$$FV^{-1} = \begin{bmatrix} \frac{\beta\lambda\theta\gamma n}{cd(\delta_1+\gamma)} & \frac{\beta\lambda\theta n}{cd} & \frac{\beta\lambda\theta}{cd} \\ 0 & 0 & 0 \\ 0 & 0 & 0 \end{bmatrix} \tag{S13}$$

And the $R_0$ of strain $i$ is given by the spectral radius of this matrix:

$$R_0 = \frac{\beta\lambda\theta\gamma n}{cd(\delta_1 + \gamma)} = \frac{\beta_{wt}(\frac{1+h}{1+he^{\Delta\Delta G_i/RT}})^K \lambda\gamma n}{cd(\delta_1 + \gamma)(1 + \frac{Ab}{K_{D,wt}} e^{-\Delta\Delta G_{Ab,i}/RT})} \tag{S14}$$

##### Landscape characterization

We measured the number of fitness optima on the receptor-binding landscape by testing whether any single amino acid substitution could increase receptor-binding affinity (decrease $\Delta\Delta G$). Only 536 variants had no neighbor with stronger receptor binding, which is remarkably low for a landscape of this size. For comparison, a maximally rugged NK landscape of the same size is expected to contain more than 33,000 local fitness optima (*62*). To determine whether this smoothness arose from the distribution of predicted binding affinities alone, we randomly permuted affinity values across genotypes. These randomized landscapes contained, on average, more than 35,000 local optima (Fig. S8), demonstrating that the smoothness arises from the organization of binding affinities in genotype space rather than their distribution alone.

We next examined how these local optima were organized in sequence space by constructing a network in which each local optimum is represented by a node and edges connect optima separated by a single mutation. The resulting graph contained four connected components, with all variants within each component having identical receptor-binding affinity and therefore forming neutral networks (Fig. 1). Some networks had non-uniform connectivity, giving genotypes substantially different fractions of neutral neighbors (Fig. S9A). However, the average fitness effect of mutations was similar across genotypes within each network (Fig. S9B), because genotypes with more neutral neighbors also tended to have more strongly deleterious non-neutral neighbors (Fig. S9C). The landscape thus exhibits a U-shaped distribution of mutant fitness effects (*63–65*), with mutations tending to be either neutral or strongly deleterious.

**Dimensionality reduction**

We projected the full quasispecies model by explicitly simulating a subset of variants. Mutational flux from tracked variants to untracked variants, calculated as the difference between the total outgoing flux from tracked variants and the incoming flux to any tracked variant, was assigned to a separate “other” compartment with its own $I_1$, $I_2$, and $V$ states, and with no back-mutation out of this compartment. The receptor- and antibody-binding affinities of the “other” compartment were set to the mean values across all untracked variants. Mutational flux into the “other” compartment was also stochastically thresholded when the expected flux was less than 1, as in the full model. We also tested a second projection method in which mutations outside the tracked set were returned to their source variants, effectively lowering the mutation rate, but this hardly changed the results.

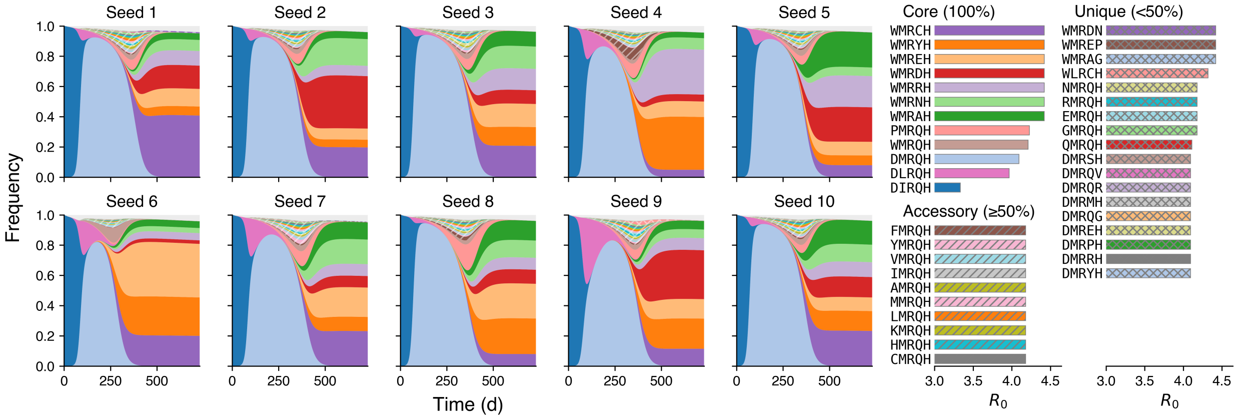


**Figure S1**: **Classification of variants captures variability across simulations.** Frequencies and $R_0$ values of the 40 most abundant variants across ten simulations. Variants are classified as core if they are among of the top 40 variants in all simulations, accessory if they are in the top 40 in at least 50% of simulations, and unique otherwise. Colors are matched with Fig. 1F.

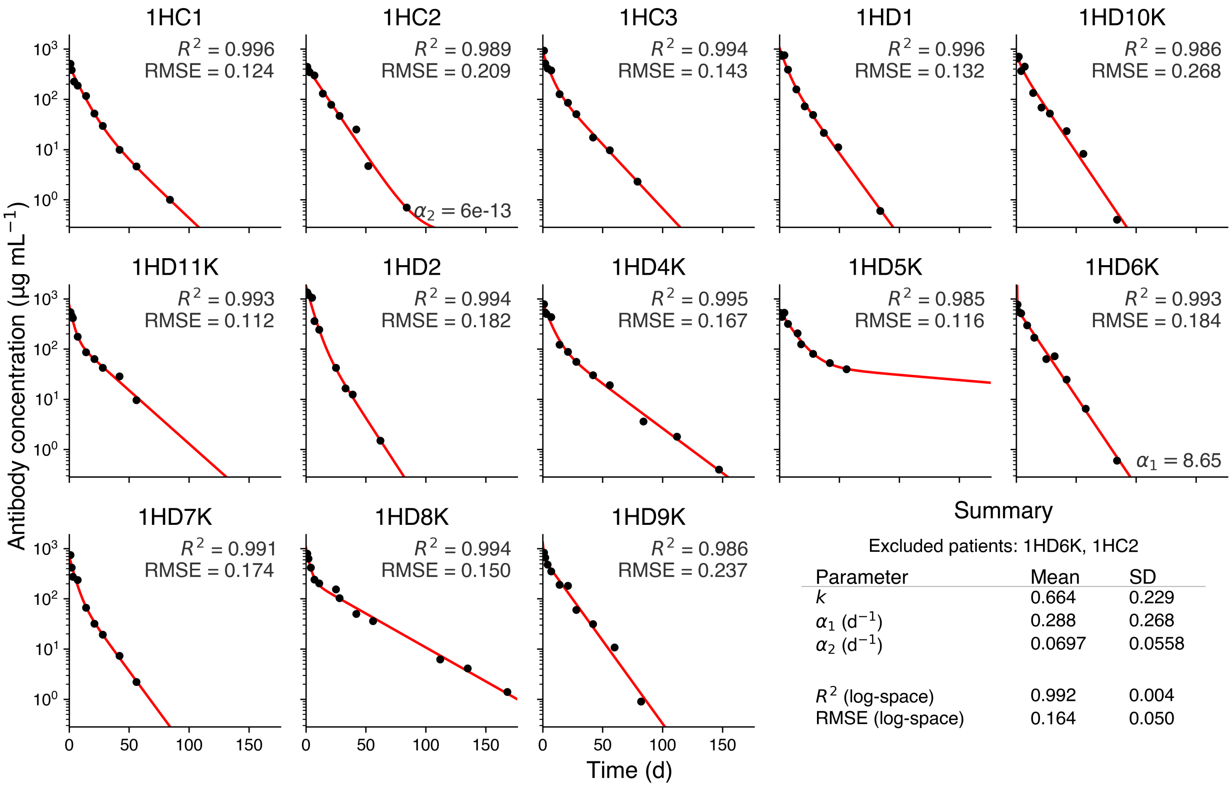


**Figure S2**: **Antibody decay after infusion of 10-1074 follows biphasic kinetics.** A biphasic decay model $[Ab](t) = [Ab]_{max}[ke^{-\alpha_1 t}+(1-k)e^{-\alpha_2 t}]$ was fit to measured serum antibody concentrations from patients given 30 mg/kg of 10-1074 in clinical trial (*12*).

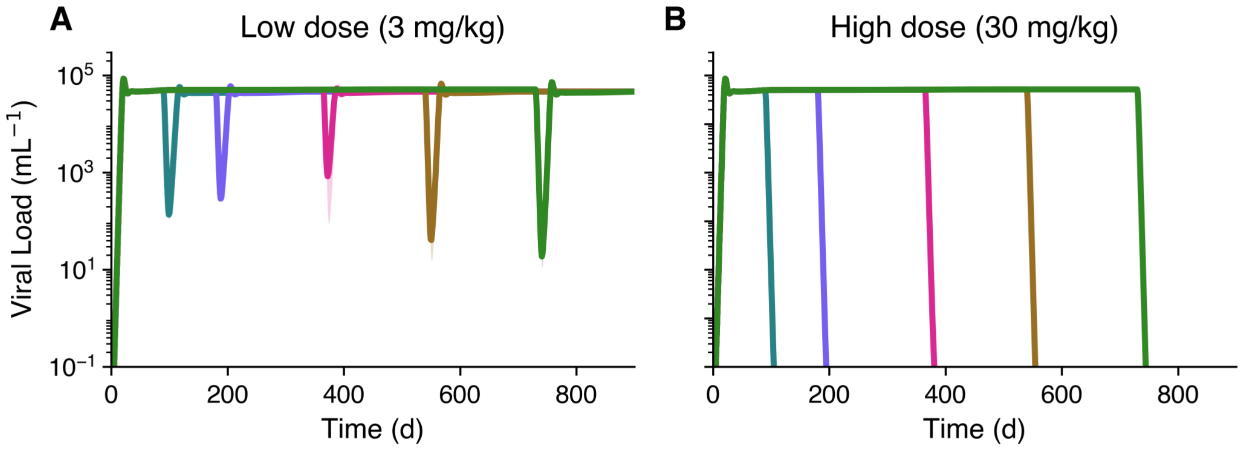


**Figure S3**: **Success of continuous 10-1074 administration is dose-dependent. (A and B)** Treatment with (A) 3 mg/kg or (B) 30 mg/kg of 10-1074 maintained at a constant concentration following administration.

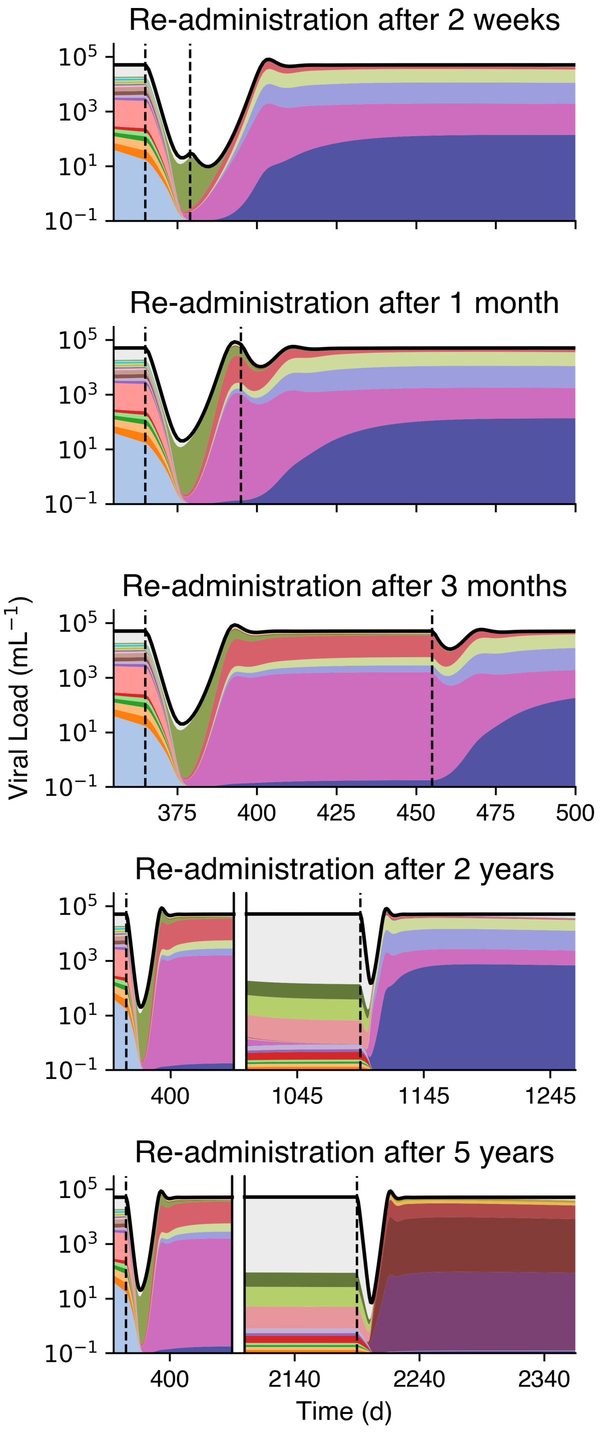


**Figure S4**: **Repeated antibody doses are less effective.** Viral load (thick line) and variant frequencies after simulated treatment with two doses of 30 mg/kg 10-1074. Variant colors are matched with Fig. 2C; the gray area represents the frequency of variants outside the most abundant set. Note that the gray area has expanded over time due to neutral drift.

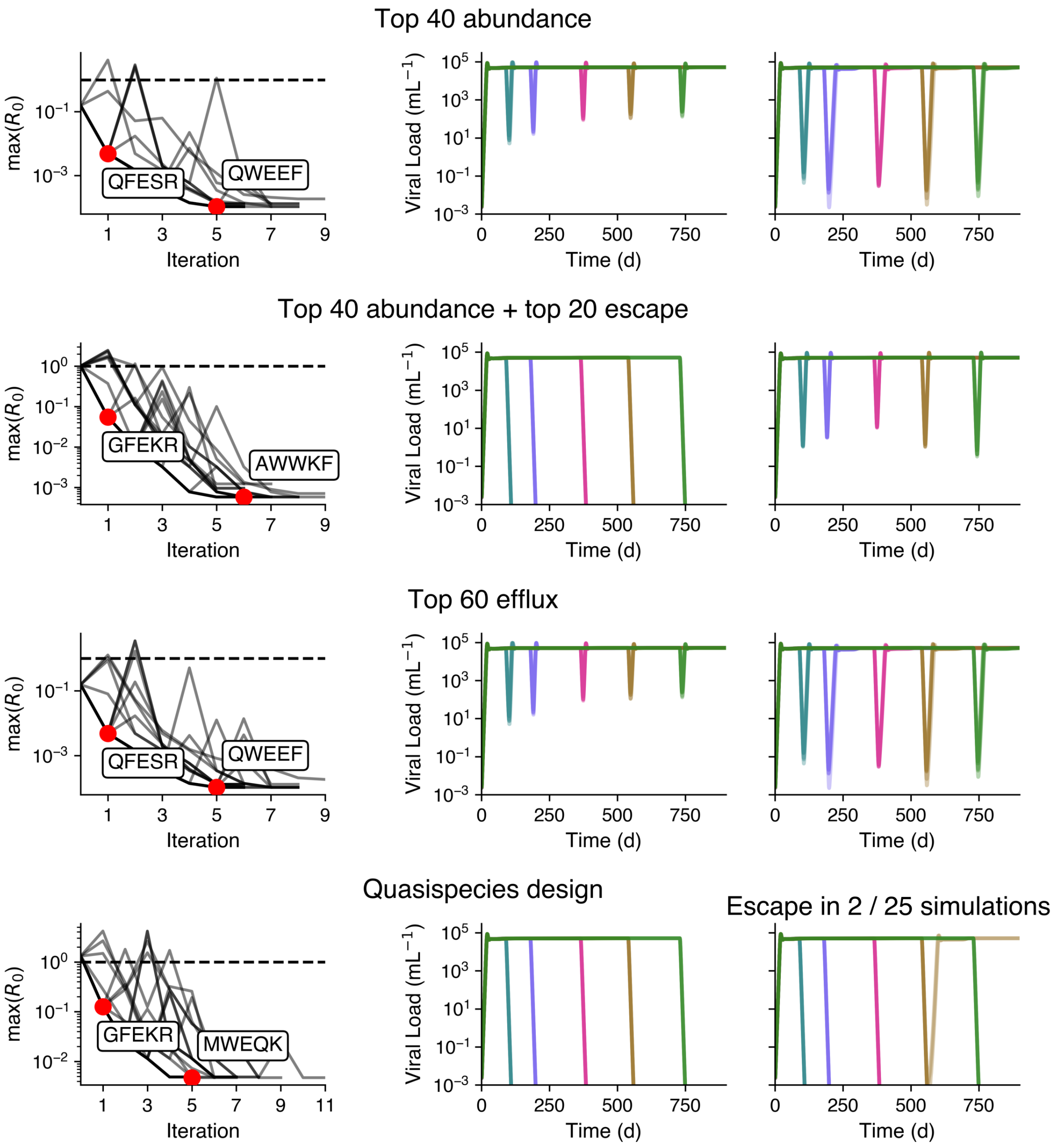


**Figure S5**: **Only quasispecies design finds multiple effective broadly neutralizing antibodies.** From left to right: trajectories from 25 optimization runs, highlighting the best antibody after a single mutation from wild type 10-1074 and the global optimum; viral loads following simulated treatment with the one-iteration optimum; and viral loads following simulated treatment with the global optimum. From top to bottom: virus panels comprise (1) the 40 most abundant variants during untreated infection; (2) these 40 variants plus the 20 must abundant escape mutants following 10-1074 administration; (3) the 60 variants with the highest mutational efflux during untreated infection; and (4) all variants with $\Delta\Delta G_{receptor} \leq 0$.

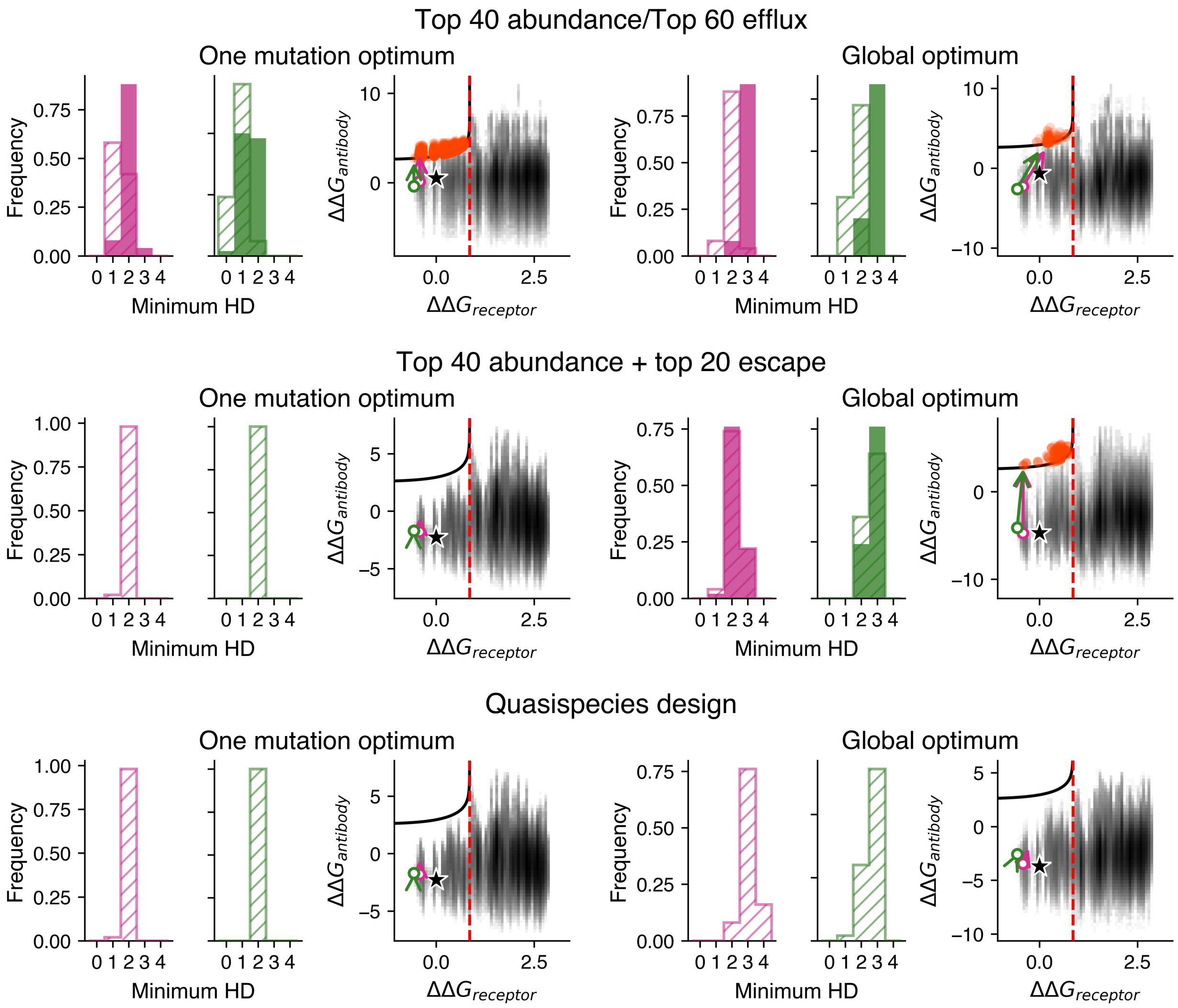


**Figure S6**: **Quasispecies design finds multiple antibodies without $R_0 > 1$ escape variants.** From left to right: minimum Hamming distance to any variant with $R_0 > 1$ (solid) or $R_0 > 0.25$ (hatched) in the presence of 30 mg/kg of designed antibody, measured for the 50 most abundant variants in the quasispecies after 1 year (purple) and 2 years (green) of untreated infection. Distribution of changes in receptor- and antibody-binding affinities for all 3.2 million mutants, with $R_0 > 1$ variants at 30 mg/kg antibody highlighted in red; Arrows denote trajectories of the mean receptor- and antibody-binding affinities prior to antibody infusion to 50 days post-treatment, as in Fig. 2D. The first three panels show results for the optimal antibody one mutation away from wild type 10-1074, and the final three panels show results for the global optimum identified across 25 optimization runs using the denoted virus panels.

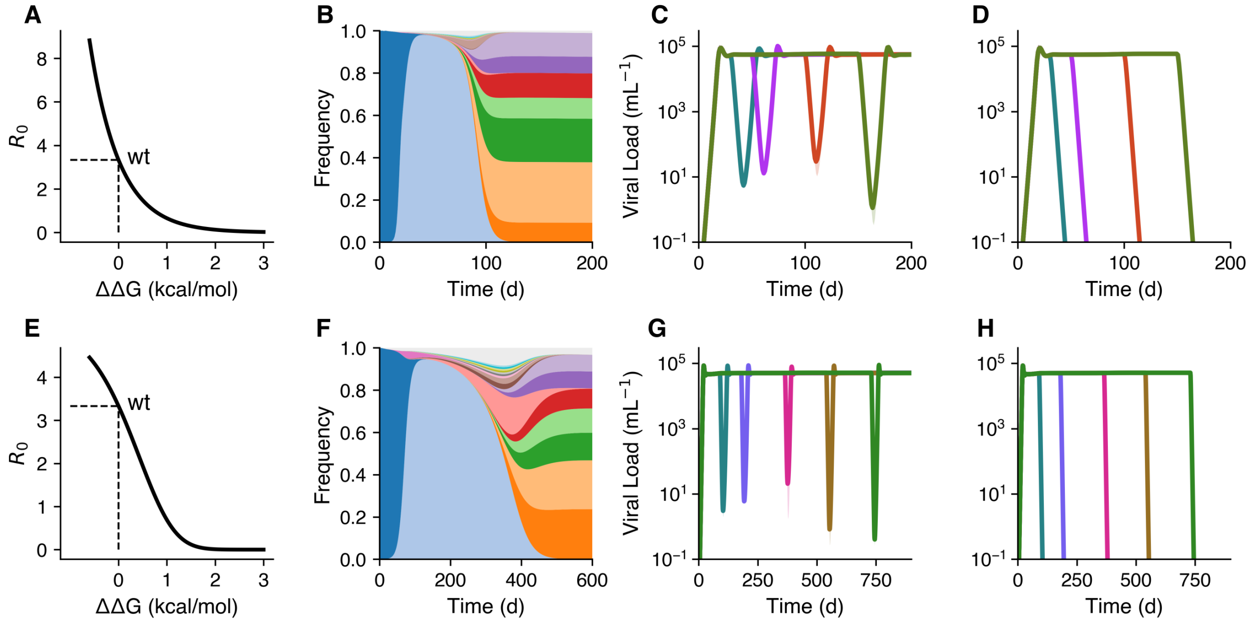


**Figure S7**: **Model results are robust to the choice of binding affinity-infection rate coupling.** **(A-D)** Main model results using the exponential dependence of Eq. S3 of the infection rate $\beta$ on receptor-binding $\Delta\Delta G$. (A) $R_0$ across relevant values of $\Delta\Delta G$ in the absence of antibody. (B) Viral abundances during untreated infection. (C and D) Viral loads following administration of (C) wild type 10-1074 or (D) the quasispecies design one-iteration optimal antibody (S96K). **(E-H)** Same as (A-D), using the Hill-like saturation for $\beta$ used in the main text (Eq. S4).

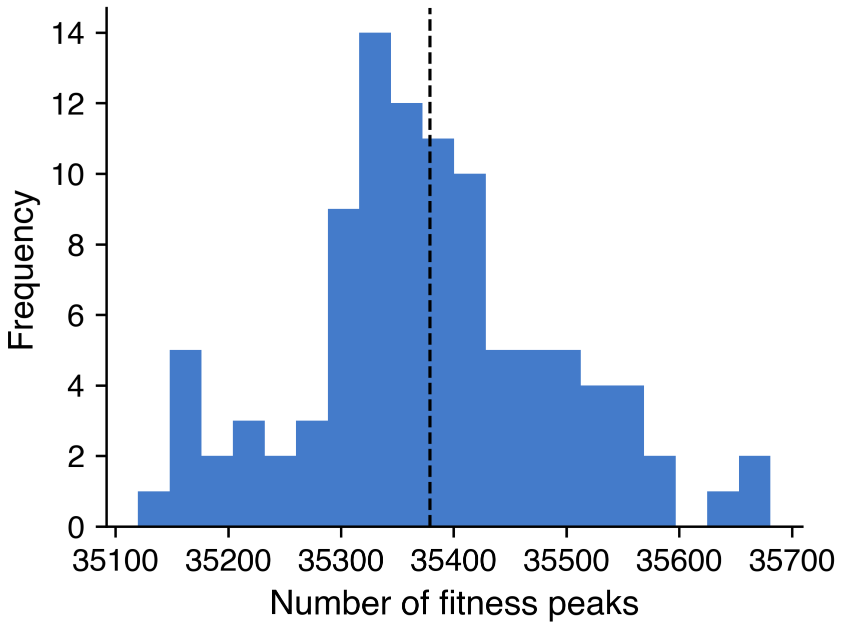


**Figure S8**: **Shuffled receptor-binding landscapes contain many more fitness peaks.** Distribution of the number of fitness peaks, defined as variants where no single mutation can increase fitness, across 100 landscapes generated by randomly assigning the observed $\Delta\Delta G$ values to genotypes. The dashed line indicates the mean across all landscapes.

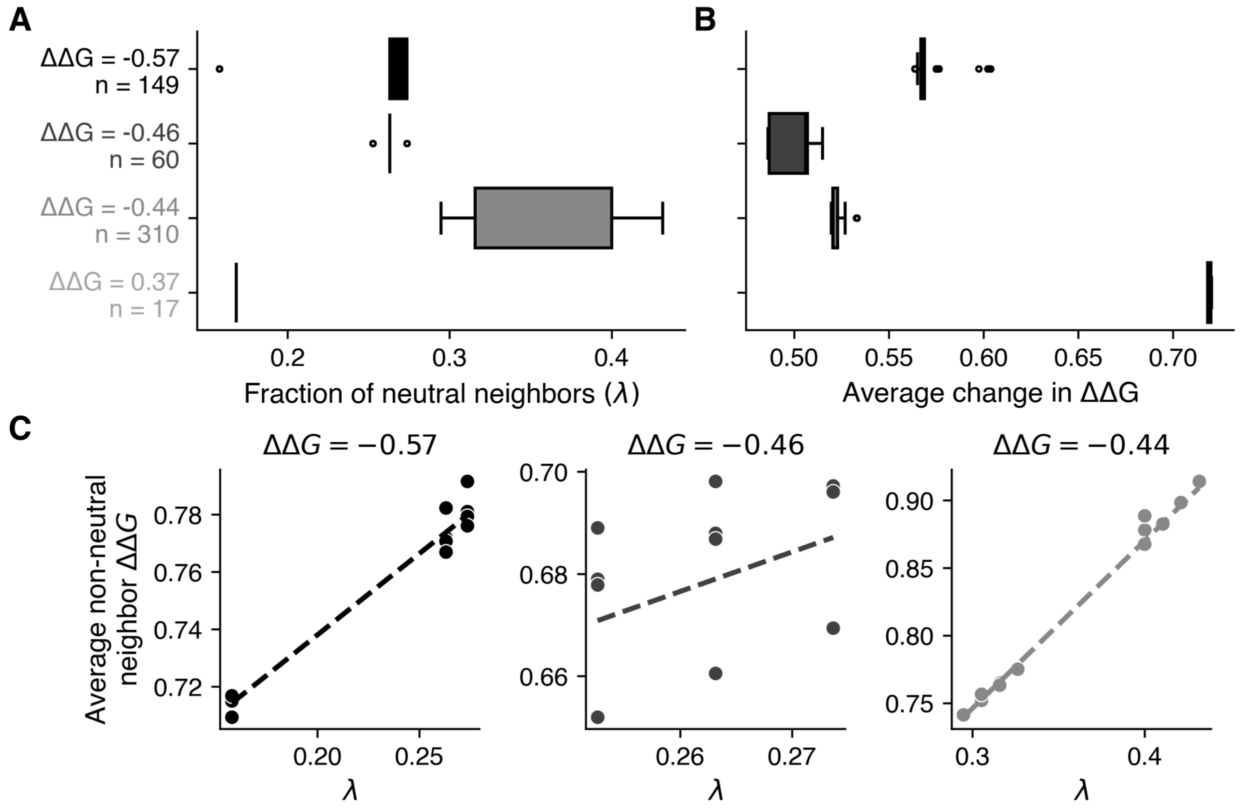


**Figure S9**: **U-shaped distribution of mutant fitness effects on neutral networks.** **(A)** Fraction of single-mutant neighbors with identical receptor-binding affinity for each variant across the four neutral networks of local optima. Box plots show the distribution across variants within each neutral network. **(B)** Distribution of the mean change in $\Delta\Delta G$ across all single-mutant neighbors for these variants. **(C)** Relationship between the fraction of neutral neighbors and the mean $\Delta\Delta G$ of non-neutral neighbors for variants on the three neutral networks with $\Delta\Delta G < 0$.

**Table S1**: **Parameters for the quasispecies model.** Values for the basic infection dynamics parameters are taken from (*66*) unless otherwise specified.

| Parameter | Interpretation | Value |
|---|---|---|
| | *Basic infection dynamics* | |
| $\lambda$ | Target cell production rate | $10^7$ cells $d^{-1}$ |
| $d$ | Target cell loss rate | 0.2 $d^{-1}$ |
| $\beta_{wt}$ | Infection rate of wild-type virus | $2 \times 10^{-9}$ virions$^{-1}$ $d^{-1}$ |
| $\gamma$ | Eclipse phase progression rate | 1 $d^{-1}$ |
| $\mu$ | Per-site mutation rate | $10^{-4}$ (*50*) |
| $\delta_1$ | Death rate of $I_1$ cells | 0.5 $d^{-1}$ |
| $\delta_2$ | Death rate of $I_2$ cells | 1 $d^{-1}$ |
| $n$ | Burst size of infected cells | 1000 |
| $c$ | Clearance rate of virus particles | 20 $d^{-1}$ |
| | *Receptor- and antibody-binding* | |
| $\Delta\Delta G_i$ | Change in receptor-binding affinity for strain $i$ | Variable |
| $\Delta\Delta G_{i,Ab}$ | Change in antibody-binding affinity for strain $i$ | Variable |
| $RT$ | Product of ideal gas constant and temperature | 0.616 kcal $mol^{-1}$ |
| $h$ | Saturation location parameter | 0.1 |
| $K$ | Saturation shape parameter | 5 |
| $K_{D,wt}$ | Dissociation constant of wild type gp120-10-1074 binding | $10^{-8}$ M (*54*) |
| | *Antibody dynamics* | |
| $[Ab]_{max}$ | Maximal antibody concentration after administration | $2.8 \times 10^{-6}$ M |
| $\alpha_1$ | Fast antibody decay rate | 0.288 $d^{-1}$ |
| $\alpha_2$ | Slow antibody decay rate | 0.070 $d^{-1}$ |
| $k$ | Proportion of fast decay | 0.664 |

**Caption for Movie S1. Antibody administration causes long-lasting reorganization of the quasispecies.** Animated version of Figure 3E showing the quasispecies response to 10-1074 administration. Nodes represent viral variants and are colored by abundance, and edges connect single-mutational neighbors and are colored by mutational flux.
